# Shigella O-specific polysaccharide functional IgA responses mediate protection against shigella infection in an endemic high-burden setting

**DOI:** 10.1101/2023.05.04.539451

**Authors:** Biana Bernshtein, Meagan Kelly, Deniz Cizmeci, Julia A. Zhiteneva, Ryan Macvicar, Mohammad Kamruzzaman, Taufiqur R. Bhuiyan, Fahima Chowdhury, Ashraful Islam Khan, Firdausi Qadri, Richelle C. Charles, Peng Xu, Pavol Kováč, Robert W. Kaminski, Galit Alter, Edward T. Ryan

**Author notes:** Co-senior (Galit Alter, Edward T. Ryan).

## Abstract

Shigella is the second leading cause of diarrheal disease-related death in young children in low and middle income countries. The mechanism of protection against shigella infection and disease in endemic areas is uncertain. While historically LPS-specific IgG titers have been associated with protection in endemic settings, emerging deeper immune approaches have recently elucidated a protective role for IpaB-specific antibody responses in a controlled human challenge model in North American volunteers. To deeply interrogate potential correlates of immunity in areas endemic for shigellosis, here we applied a systems approach to analyze the serological response to shigella across endemic and non-endemic populations. Additionally, we analyzed shigella-specific antibody responses over time in the context of endemic resistance or breakthrough infections in a high shigella burden location. Individuals with endemic exposure to shigella possessed broad and functional antibody responses across both glycolipid and protein antigens compared to individuals from non-endemic regions. In high shigella burden settings, elevated levels of OSP-specific FcαR binding antibodies were associated with resistance to shigellosis. OSP-specific FcαR binding IgA found in resistant individuals activated bactericidal neutrophil functions including phagocytosis, degranulation and reactive oxygen species production. Moreover, IgA depletion from resistant serum significantly reduced binding of OSP-specific antibodies to FcαR and antibody mediated activation of neutrophils and monocytes. Overall, our findings suggest that OSP-specific functional IgA responses contribute to protective immunity against shigella infection in high-burden settings. These findings will assist in the development and evaluation of shigella vaccines.

## Introduction

Diarrheal diseases caused by enteric pathogens are a major cause of death in children younger than 5 years of age in low and middle income countries (LMIC)(*1, 2*). Attempts to develop effective vaccines against enteric pathogens have been impeded by both the breadth of pathogens involved, and often relative cursory understanding of both correlates and mediators of protective immunity against individual pathogens in endemic settings. Vaccines that have been protective against enteric pathogens in non-endemic settings have sometimes failed or provided decreased protection when evaluated in endemic areas. For example, licensed rotavirus vaccines that are extremely effective in resource-rich countries have shown reduced efficacy in LMIC settings(*3*). Additionally, an oral live attenuated shigella vaccine that was promising in North American vaccinees was not immunogenic in Bangladeshis(*4*), and an oral live attenuated cholera vaccine failed in an endemic zone field study after showing excellent protection in a North American experimental infection-based evaluation(*5*). Deciphering correlates of protection against enteric infection in endemic and non-endemic areas is key to informing efficacious vaccine development for use in endemic areas.

Shigella infection accounts for tens of thousands of deaths each year and is the second leading diarrheal-related cause of death in young children(*2*). Four pathogenic bacterial species cause shigellosis, defined by their O-specific polysaccharide (OSP) - *Shigella flexneri, S. sonnei, S. boydii* and *S. dysenteriae*. The majority of shigellosis in low resource settings is caused by *S. flexneri*, while *S. sonnei* causes shigellosis in industrialized and transitioning countries(*6*). *S. flexneri* exists as a number of serotypes, with *S. flexneri* 2a, 3a and 6 being among the most common globally(*6*). A number of correlates of protection against shigellosis have been proposed. LPS-specific IgG protects against natural shigella infection in a serotype-specific manner(*7*–*9*), while LPS-specific IgA is induced in response to infection in shigella endemic regions (*10, 11*). In a controlled human challenge of North American individuals with *S. flexneri* 2a, pre-existing functional antibodies that recognized IpaB predicted protection against severe shigellosis(*12*), while in a separate *S. sonnei* challenge study in which North American individuals were pre-screened and excluded for *S. sonnei* LPS-specific IgG (but not IgA) in their serum(*13*), shigella LPS-specific IgA predicted protection from shigellosis following challenge. These data suggest that LPS-specific IgA, IgG and IpaB-specific functional antibodies have the potential to protect against shigellosis, however the mechanism(s) of protection is unclear. We thus applied a systems approach to compare the shigella-specific antibody responses in individuals living in shigella endemic and non-endemic areas, and to identify correlates of protection in a shigella-endemic location.

## Results

### Shigella-specific antibody profiles are distinct between individuals living in endemic and non-endemic settings

Individuals living in shigella endemic regions are exposed to shigella infection from childhood and develop partially protective immunity against shigellosis during the first years of life(*14*). Yet, our understanding of the humoral immune response that evolves in these individuals is incomplete. To this end, we compared shigella-specific antibody landscapes found in individuals living in a shigella—endemic area in Peru (endemic) to those in individuals living in North America (non-endemic) to evaluate the extent of the difference in shigella--specific humoral immune responses. Specifically, we analyzed serum samples from a subset of military recruits in Peru upon entry to a training camp (*15*) (n=55) and a subset of individuals participating in a separate *S. sonnei* 53G U.S. shigella challenge study at the pre-challenge time point(*16*) (n=44). Individuals participating in the challenge study were prescreened for low shigella LPS-specific IgG, as a marker for lack of previous exposure. Shigella-specific antibody isotype, subclass, and Fc-receptor (FcR) binding profiles across IpaB, IpaC, IpaD, LPS from *S. flexneri* 2a, 3a, 6 and *S. sonnei*, and OSP from *S. flexneri* 2a, 3a and *S. sonnei* were analyzed. To compare endemic and non-endemic shigella-specific humoral profiles, a Least Absolute Shrinkage and Selection Operator (LASSO) was used to down select a set of minimal antibody features that differed most between the endemic and non-endemic groups (**Fig1A**). A Partial Least Squares-Discriminant Analysis (PLS-DA) was used to visualize the data (**Fig1A**). As few as 8 out of total 99 analyzed antibody-features captured by systems serology were sufficient to separate individuals living in endemic and non-endemic regions. While samples from non-endemic regions were enriched with IgM and IgA2 antibodies against IpaC, antibodies of individuals living in shigella-endemic regions harbored a mature antibody response underlined by expanded FcR binding of IpaC, IpaD and OSP-specific antibodies. To define a minimal set of biomarkers associated with each group, the LASSO algorithm eliminates co-correlated features and selects a minimal set of features that accounts for variation across individuals. The co-correlates of the LASSO-selected features may help identify overall antibody profiles that differ most in the response to shigella across individuals living in endemic and non-endemic regions. LASSO-feature correlation network analyses revealed 3 correlation networks (r>0.85, q<0.01) (**Fig1B**). FcγR3b binding of IpaD specific antibodies and FcγR2a binding of IpaC-specific antibodies correlated with each other and with IpaB, OSP and LPS IgG1 and FcγR binding, underlining the abundant and broad shigella-specific class switched IgG functional antibody response in individuals living in endemic settings. IgM specific to *S. sonnei* OSP were enriched in non-endemic settings, and correlated with IgM specific to IpaB, IpaC and *S. sonnei* LPS, suggesting a non-class switched immune response is differentially enriched in non-endemically exposed individuals to both protein and glycolipid targets. Importantly, *S. sonnei* OSP-specific IgA1 responses correlated with *S. sonnei* LPS-specific IgA1 levels, as these two antigens share the same OSP. IpaC, IpaB and *S. flexneri 2a* and *S. sonnei* OSP-specific IgG1 and Fcγr2a, 3a, 3b and FcαR binding antibodies were significantly enriched in individuals living in endemic regions (**Fig1C**). Collectively, these data suggest that individuals who live in a shigella-endemic region harbor a more IgG class-switched, mature and broad shigella-specific antibody response. Antibodies specific to both shigella proteins and LPS/OSP found in their serum bind to multiple FcRs and harbor the potential to activate innate immune cells.

**Fig 1.**
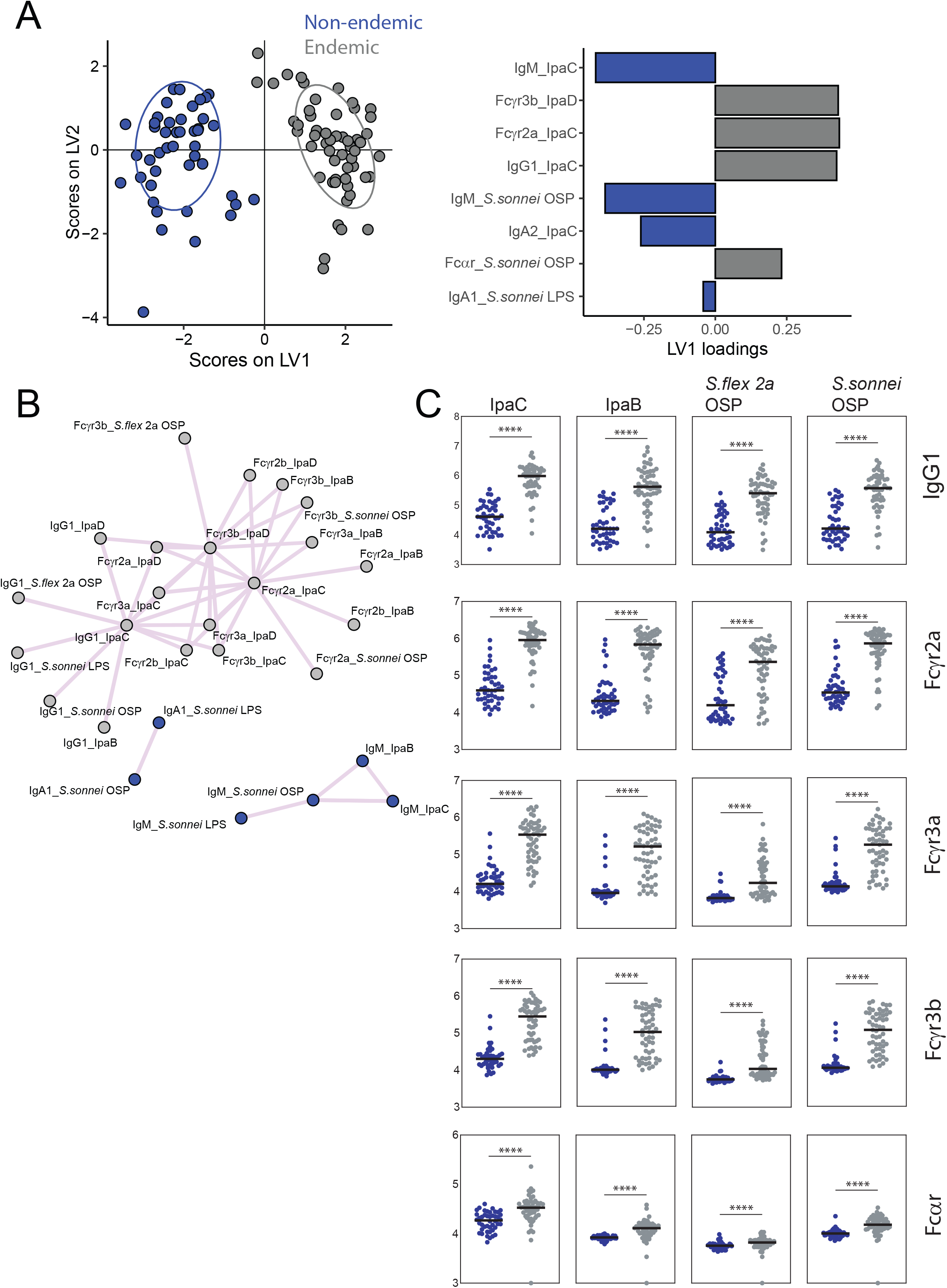
Enrichment of FcR binding shigella-specific antibodies in individuals living in shigella endemic region. **(A)** Score plots and LV1 loadings of PLS-DA model using LASSO selected features of shigella-specific antibody isotypes and FcR binding in serum of individuals living in shigella endemic or non-endemic regions. **(B)** Correlation network of antibody features presented in A (r>0.85, p corrected p<0.01) **(C)** Shigella -specific IgG1 and binding to Fcγr2a, Fcγr3a, Fcγr3b and FcαR in serum of individuals living in shigella endemic or non-endemic regions. (non-endemic n=44, endemic n=54) (*p<0.05, **p<0.01, ***p<0.001, ****p<0.0001, Mann-Whitney test)

### *Shigella sonnei* challenge induces functional shigella-specific antibodies in North American individuals

In endemic settings, exposure to shigella occurs early in life, and by adulthood most individuals are partially protected from shigellosis(*14, 17*). We sought to determine whether a broad and mature shigella-specific humoral immune response could develop in adults previously unexposed to shigella infection, upon challenge with *S. sonnei*. Previously, we have demonstrated that controlled *S. flexneri* 2a challenge induces FcR binding antibodies in healthy North American volunteers(*12*). Using a systems approach we analyzed serum samples of individuals participating in a controlled *S. sonnei* challenge study at baseline, day 14, 28 and 56 post-challenge (**Fig2A**). We measured shigella-specific antibody isotype, subclass, FcR binding as described above, and OSP-, IpaB- and IpaD-specific antibody mediated complement deposition (ADCD) and neutrophil phagocytosis (ADNP). Study participants were pre-screened for *S. sonnei* LPS specific IgG, yet some individuals had increased levels of *S. sonnei* OSP-specific IgA(*13*). Individuals that harbored increased levels of OSP-specific IgA1 were protected from shigellosis(*13*)(**Fig2B**); however, OSP-specific IgA2 and IgG1 were not augmented in these protected individuals. (**Fig2B**). Although antibody-mediated complement deposition is often used as a measure of antibody functionality and protection against bacterial infection(*18*), baseline levels of complement deposition mediated by *S. sonnei* OSP-specific antibodies did not correlate with protection (**Fig2B**). Finally, we investigated the humoral evolution of IpaC, IpaB, *S. flexneri* 2a OSP and *S. sonnei* OSP-specific antibodies at days 14, 28 and 56 following challenge (**Fig2C**). IpaC, and more pronouncedly IpaB-specific FcR binding antibodies increased rapidly following challenge. IpaB-specific IgG1 remained high at 56 days post challenge, as well as binding of IpaB antibodies to FcγR3b (**Fig2C**). OSP-specific antibodies appeared quickly after challenge; however, only *S*.*sonnei* OSP-specific antibodies exhibited increased binding to FcγRs over time, pointing to development of a species-specific functional antibody response. Binding to FcαR was-specifically robust and pronounced for *S. sonnei* OSP-specific antibodies post-challenge (**Fig2C**). Overall, controlled *S. sonnei* challenge of North American healthy adults induced a robust and functional shigella-specific antibody response.

**Fig 2.**
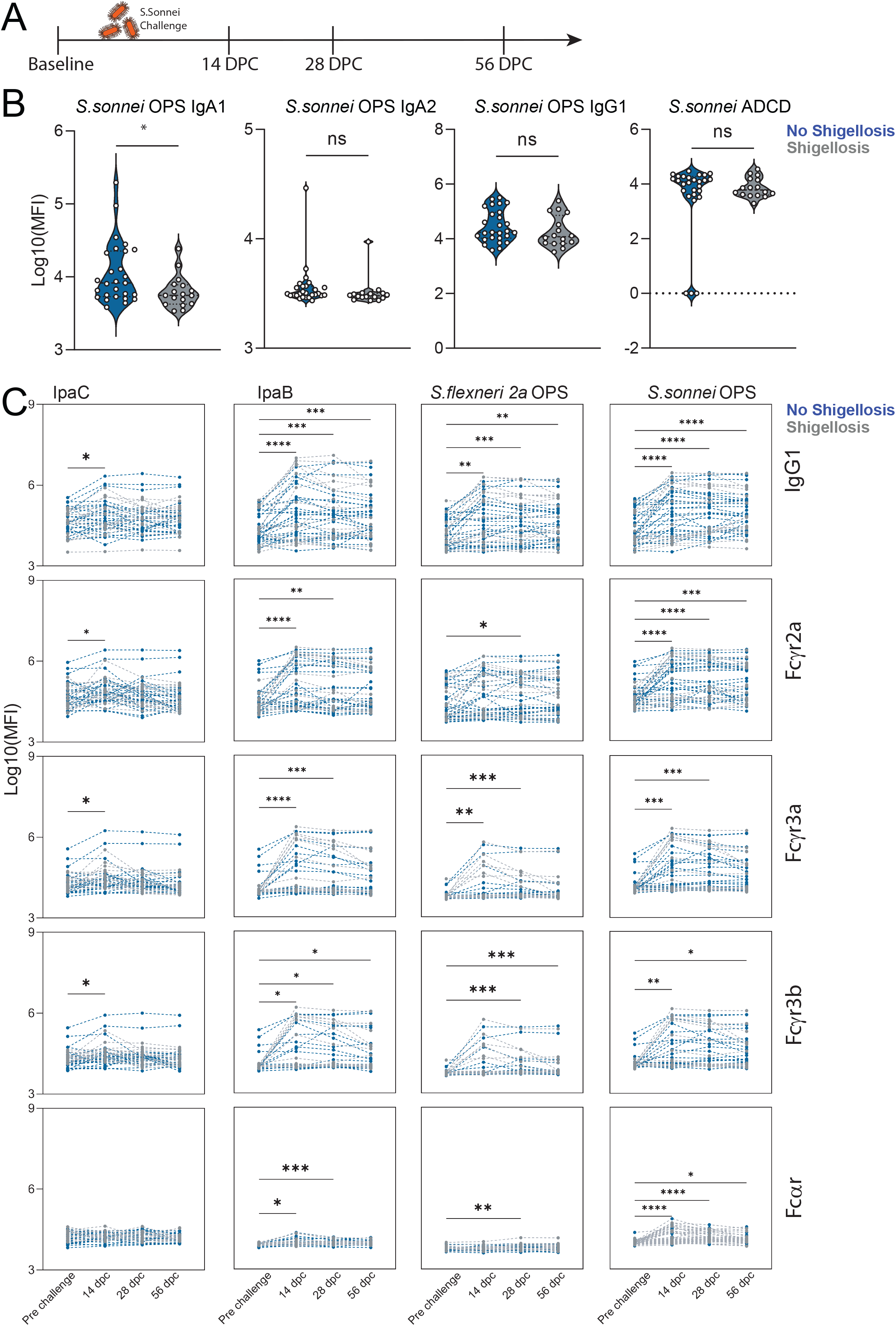
Challenge induced antibody features. **(A)** *S. sonnei* challenge timeline. **(B)** *S. sonnei* OSP-specific IgA1, IgA2 and IgG1 levels and antibody dependent complement deposition (ADCD) in serum of challenged individuals at baseline. (n=44) *p<0.05 **(C)** Humoral evolution of IpaC, IpaB, *S. flexneri* 2a OSP and *S. sonnei* OSP-specific IgG1 and binding to Fcγr2a, Fcγr3a, Fcγr3b and FcαR in serum of individuals pre and post challenge with *S. sonnei*. (n=44) (*p<0.05, **p<0.01, ***p<0.001, ****p<0.0001, Wilcoxon test)

### Shigella OPS-specific antibodies protect against infection in high shigella burden settings

The shigella-specific antibody profile in Peruvian army recruits at entry to camp was broad and mature. Yet, the exposure risk to shigellosis in this setting was extremely high with a diarrheal attack rate of 32% and *Shigella* spp cultured in ∼40% of the diarrheal stools(*15*). For this analysis, a subset of the recruits that developed shigellosis were included, albeit infection occurred at different time points across the cohort. To define whether similar or distinct antibody profiles were associated with shigellosis in endemically exposed individuals, we analyzed longitudinal serum samples from a set (n=34) of individuals with a known *S. flexneri* 2a infection over the study period. We divided the samples into two groups: susceptible – infected within 90 days of entering the camp (n=26) or resistant – infected later than 90 days after entering the camp (n=8). Using systems serology, we analyzed shigella-specific antibody isotype, subclass, FcR binding, ADCD and ADNP. To determine the differences in shigella-specific antibody profiles between resistant and susceptible groups, a LASSO was used to down select a set of minimal antibody features that differed most between two groups and PLS-DA was used to visualize the data (**Fig3A**). Just 3 antibody features, out of 99 profiled per sample, were sufficient to discriminate between individuals that were susceptible and resistant in the first 3 months of follow-up, including FcαR binding of *S. flexneri* 2a OSP-specific antibodies, *S. flexneri* 2a OSP-specific ADNP and *S. sonnei*-specific LPS IgM, all features were significantly increased in resistant individuals (**Fig3B**). Though complement deposition has been suggested to mediate protection against shigellosis, resistant individuals did not show augmented *S. flexneri* 2a OSP-specific ADCD (**Fig3B**). Importantly, both IgG and IgA can drive neutrophil activation, due to the constitutive expression of both FcγR3b and FcαR on the surface of neutrophils(*19, 20*). Thus, given that *S. flexneri* 2a OSP-specific antibodies displayed increased FcαR binding and ADNP, we sought to investigate whether OSP-specific IgA (that interact with FcαR) mediates neutrophil phagocytosis, potentially by binding to FcαR expressed by neutrophils. To address the role of OSP-specific IgA in resistant individuals, we depleted IgA from pools of sera from shigella susceptible and resistant cohorts and performed *S. flexneri* 2a OSP-specific ADNP assays with depleted and non-depleted serum (**Fig3C**). IgA depletion significantly decreased *S. flexneri* 2a OSP-specific ADNP in resistant, but not susceptible individuals (**Fig3C**). Overall, our data suggests that OSP-specific IgA mediates protective neutrophil phagocytosis in shigellosis resistant individuals. To further investigate the neutrophil bactericidal activity of OSP-specific IgA, we depleted IgA from eight shigella resistant serum samples collected at entry to camp. Depletion of IgA, and consequent loss of FcαR binding, was verified by Luminex analysis (**Fig4A**). IgA depletion significantly reduced OSP-specific ADNP and antibody dependent monocyte phagocytosis (ADMP) of shigella resistant individuals (**Fig4B**). Moreover, IgA depletion reduced neutrophil secretion of primary (MPO), secondary (lactoferrin) and tertiary (MMP9) granules in response to *S. flexneri* 2a OSP-specific antibodies (**Fig4C**). Finally, reactive oxygen species (ROS) generation by neutrophils was observed in the presence of OSP-specific IgA, and significantly reduced by IgA depletion (**Fig4D**). Altogether, OSP specific IgA in shigella-resistant individuals are key to the activation of neutrophil phagocytosis and degranulation, resulting in secretion of bactericidal granules and production of ROS.

**Fig 3.**
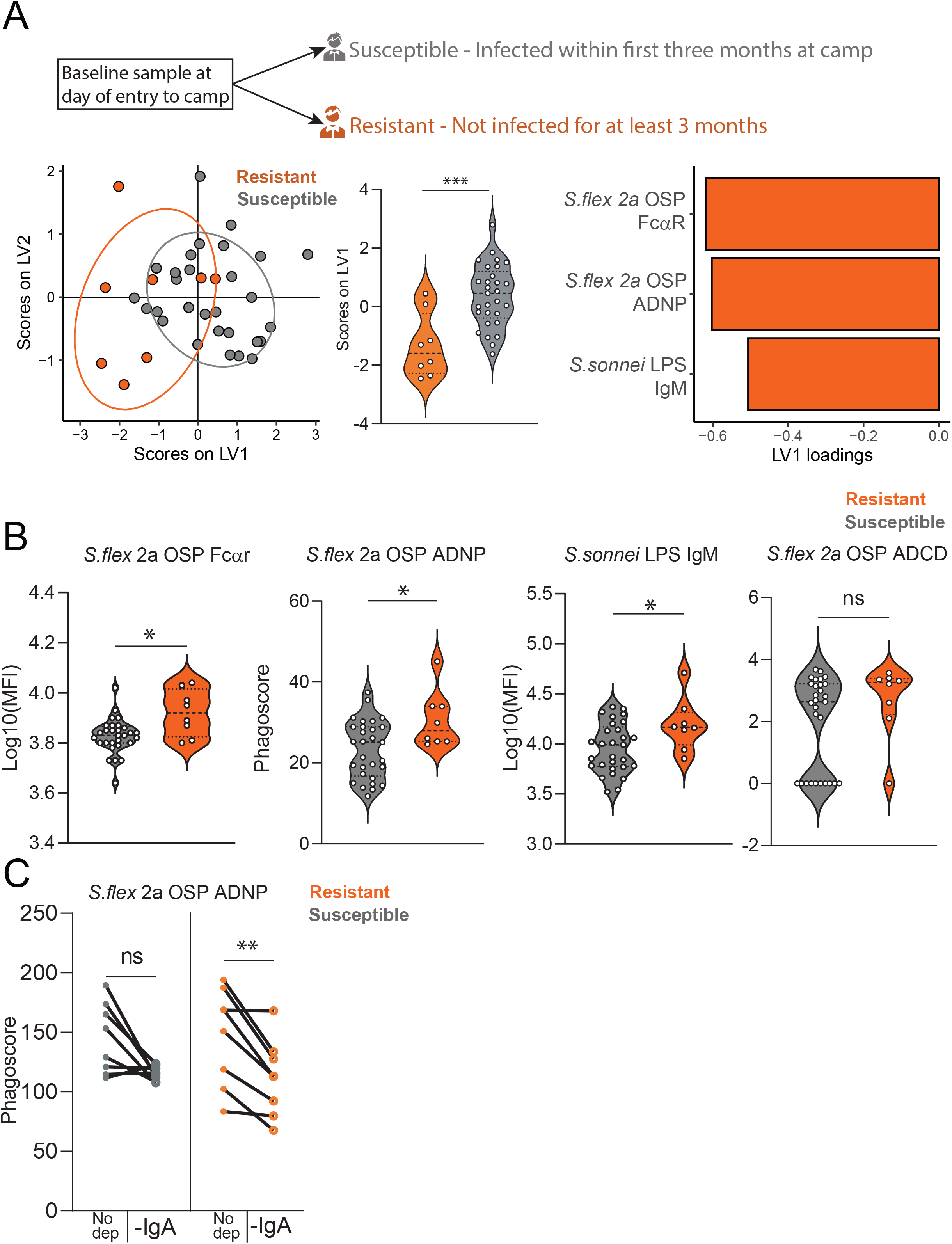
Correlates of protection against shigellosis in shigella high burden settings. **(A)** Score plots and LV1 loadings of PLS-DA model using LASSO selected features of shigella - specific antibody isotypes and FcR binding in serum of individuals susceptible (n=29) or resistant (n=8) to shigellosis at entry to camp. (***p<0.001, Mann-whitney test) **(B)** *S. flexneri* 2a OSP-specific FcaR binding and Antibody dependent neutrophil phagocytosis (ADNP), *S. sonnei* OSP-specific IgM and *S. flexneri* 2a OSP ADCD in shigellosis resistant and susceptible individuals at entry to camp. **(C)** *S. flexneri* 2a OSP ADNP in IgA depleted or non-depleted pools of serum of shigellosis resistant and susceptible individuals at entry to camp.

**Fig 4.**
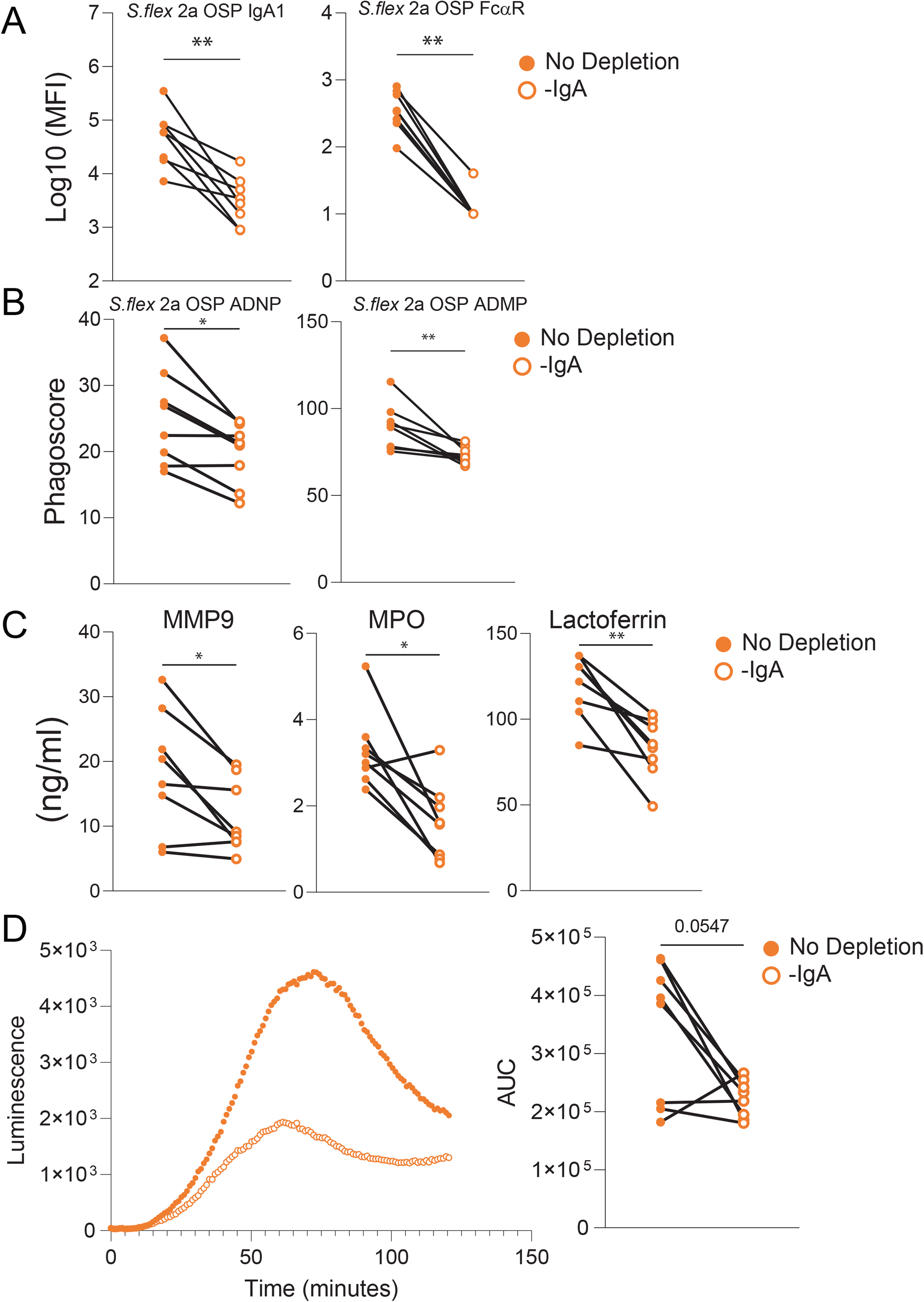
Functional OSP-specific IgA of shigellosis resistant individuals activates neutrophils and monocytes. (**A)** *S. flexneri* 2a OSP-specific IgA1 and FcαR binding in IgA depleted or non-depleted serum of shigellosis resistant individuals at entry to camp. **(B)** *S. flexneri* 2a OSP ADNP and ADMP in IgA depleted or non-depleted serum of shigellosis resistant individuals at entry to camp. **(C)** Degranulation of neutrophils in response to incubation with *S. flexneri* 2a OSP -specific immune complexes of IgA depleted or non-depleted serum of shigellosis resistant individuals. Primary (MPO), secondary (lactoferrin) or tertiary (MMP9) granules were measured by ELISA. **(D)** Reactive oxygen species luminescence of neutrophils incubated with *S. flexneri* 2a OSP-specific immune complexes of IgA depleted or non-depleted serum of shigellosis resistant individuals. AUC = area under curve. (*p<0.05, **p<0.01, Wilcoxon test)

Although shigella-resistant individuals were not infected during the first 3 months of follow-up following entry into the camp, they all eventually became infected at a later time point. We hypothesized that if OSP-specific FcαR binding functional antibodies do play a critical role in protection against shigellosis, that in a person who eventually became infected that these immune responses would decline before infection. To test this, we compared levels of FcαR-binding *S. flexneri* 2a OSP-specific antibodies at entry to the camp (baseline) and at the last collection timepoint prior to infection (pre-infection) in the 8 initially resistant individuals (**Fig5A**). Corroborating the protective role of these antibodies, the level of OSP-specific FcαR binding antibodies significantly declined over time prior to eventual infection in these individuals (**Fig5A**). Importantly, reduced *S. flexneri 2a* OSP-specific FcαR binding antibodies were not a result of a general decline in shigella-specific FcαR binding antibodies in these individuals, as FcαR binding levels of *S. flexneri* 3a OSP, *S. sonnei* OSP and IpaB-specific antibodies remained unaltered (**Fig5B**). However, this loss of *S. flexneri 2a* OSP-specific FcαR-binding was linked to a loss of ADNP and ADMP activity over time in the camp (**Fig5C**), suggesting that OSP-specific FcαR-binding antibodies that mediate ADNP and ADMP represent a functional correlate of protection against shigellosis in high shigella burden settings.

**Fig 5.**
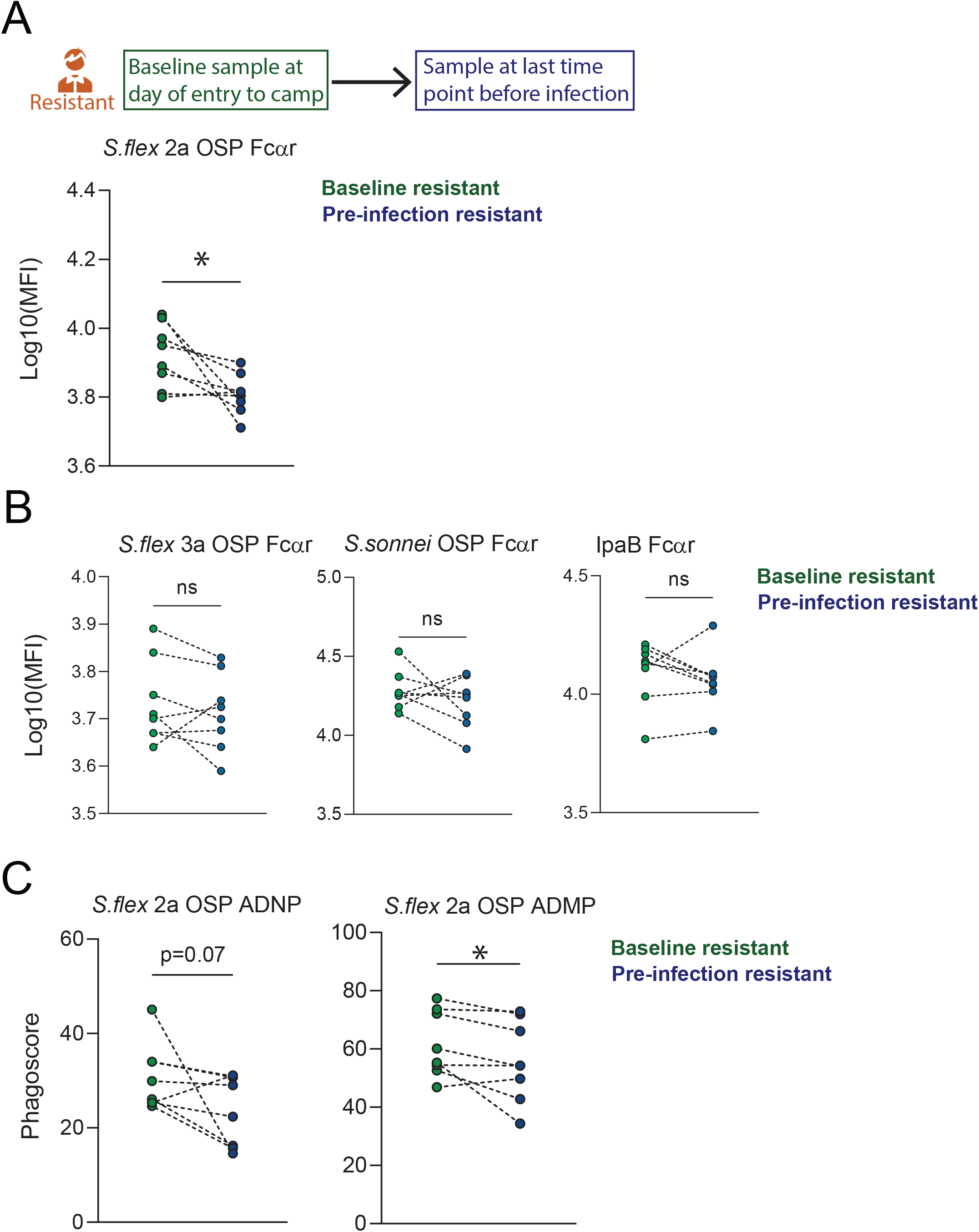
OSP-specific antibody functionality is strain-specific and is reduced prior to infection. **(A)** *S. flexneri* 2a OSP-specific FcαR biding at entry to camp and pre-infection timepoint **(B)** *S. flexneri* 3a, *S. sonnei* OSP and IpaB-specific FcαR binding at entry to camp and pre-infection timepoint **(C)** *S. flexneri* 2a OSP ADNP and antibody dependent monocyte phagocytosis (ADMP) at entry to camp and pre-infection timepoint.

## Discussion

Studies conducted decades ago in the Israeli army established that antibodies that protect against shigellosis largely target O-polysaccharides that are species and serotype-specific, suggesting that a multi-valent vaccine may be required to provide meaningful protection against shigellosis(*7, 21, 22*). This immunity can be augmented to some degree by responses targeting antigens conserved across *Shigella* spp, including invasion protein antigens (Ipa). Indeed we have recently shown, using samples collected in a *S. flexneri* 2a North American challenge study, that IpaB-specific IgG that binds to Fcγ receptors and activates monocyte and neutrophil phagocytosis correlate with protection against shigellosis(*12*). These various observations prompted us to expand our approach to study functional antibody mediated protection against shigellosis in endemic settings, where the largest burden of shigellosis is borne. First, we found marked differences in baseline shigella-specific antibody profiles when comparing individuals living in endemic and non-endemic settings. Namely, shigella-specific antibody profiles of Peruvian (endemic) individuals were marked by class-switched and FcR binding antibodies against both protein and polysaccharide shigella antigens, compared to North American (non-endemic) individuals. These immune responses in Peruvian recruits would be consistent with the previous and repetitive exposure to *Shigella* spp in this region. Second, we observed that North American volunteers challenged with wild type *S. sonnei* organisms were able to develop broad anti-shigella immune responses including antibodies that bind to Fc receptors, similar to what we previously observed in North American volunteers challenged with wild type *S. flexneri* 2a(*12*). While these findings in North American adults may be relevant to future development of traveler vaccines, neither of these observations addressed identification of potential correlates of protection against shigellosis in endemic areas.

Identifying a clear correlate of protection against shigella infection in endemic settings is impeded by the presence of a broad pre-existing shigella-specific antibody responses resulting from ongoing and repetitive exposure to *Shigella* spp. We thus took advantage of a unique longitudinal serum sample collection from a high shigella burden setting and identified susceptible (infected within first 3 months at camp) and resistant (infected later than 3 months at camp) individuals. Resistant individuals harbored elevated levels of *S. flexneri* 2a OSP-specific FcαR binding antibodies that engaged neutrophils and induced protective cellular phagocytosis. Importantly, OSP-specific IgA levels were not elevated in resistant individuals, nor were Fcγ or IgG OSP-specific antibody responses. *Shigella* spp are invasive mucosal pathogens that spread along intestinal epithelia cells, and can infect polymorphonuclear cells, macrophages, T cells and B cells, in addition to intestinal epithelial cells.(*23*–*26*). Functional antibody and antibody-mediated activity might thus play a critical role in mediating protection against shigellosis at the mucosal surface. FcαR binds IgA and is expressed on diverse innate immune cells including neutrophils, monocytes and dendritic cells(*27, 28*). Binding of FcαR by IgA on the surface of innate immune cells results in a range of cellular functions including phagocytosis(*29*), NETosis(*30*), cytokine secretion and metabolic reprogramming(*31*). A previous evaluation of rectal biopsy tissues during shigellosis in the 1990s in Bangladesh disclosed increased presence and activation of polymorphonuclear cells, macrophages and intraepithelial T cells in infected rectal tissues, and the presence of cytokines including IL-1a, IL-1b, IFN-gamma, TNF-a, IL-1Ra, IL-6, IL-8, IL-4, IL-10 and TNF-b(*32, 33*).

Our analysis identified OSP-specific FcαR binding antibodies but not total OSP-specific IgA to be associated with protection, suggesting that qualitative rather than quantitative differences may be important. For example, differential Fc-glycosylation of IgA might explain changes in binding to FcaR. Though it has been suggested that IgA-FcaR binding interfaces are not altered by glycosylation(*34, 35*), recent studies suggest that glycosylation of IgA1 and IgA2 is distinct and can affect binding to FcαR and antibody mediated neutrophil activation (*36*). Future studies should thus address glycosylation patterns of shigella OSP-specific IgA in endemic settings to address this possibility. An important observation in our analysis is that the *S. flexneri* 2a OSP-specific FcαR binding antibodies decreased over the study period to the extent that initially protected individuals eventually became infected. This could suggest that these immune responses might be relatively short-lived (and thus might hinder successful development of protective durable immune responses from vaccination) or might to some degree reflect the intense infectious pressure that was present in this hyper-endemic setting.

Our study has a number of limitations. It was limited by the human serum samples that were available to us, both in the geographic location and numbers. It will be essential to broaden our findings to other endemic areas, including in sub-Saharan Africa and Asia. The group of shigella-resistant individuals in our study was also small (n=8), as we sought to study natural infection, preventing us from controlling for group sizes. Nevertheless, we found signatures of protection in this relatively small group that were significant when compared to a larger group (n=26) of susceptible individuals, and when analyzing serum of resistant individuals over time. Another limitation of our study is that our blood samples were restricted to antibodies in the peripheral circulation. Future analysis should include mucosal and cellular-based analyses. Finally, we did not have sufficient serum sample volume to analyze glycosylation of OPS-specific IgA and plan to address this in future studies.

Our results add to a growing body of work attempting to identify correlates and mechanisms of protection against shigellosis in endemic settings(*9, 37, 38*) and support a central role of immune responses targeting the OSP component of *Shigella* spp LPS. Our results suggest that OSP-specific FcαR binding antibodies are a potential bio-marker of protection against shigella infection in endemic high-burden settings. We found a potential mechanism of protection against shigellosis at the mucosal surface since these responses induced neutrophil phagocytosis, degranulation and ROS production. Importantly, we did not find an association of antibody mediated complement deposition with protection in our analysis. This is of note since serum (cell free) bactericidal assays that assess complement binding and activation are being used to evaluate shigella vaccine candidates. Our current study using samples from an endemic zone, and our previous analysis of samples from North American volunteers suggests that an optimal candidate vaccine against shigellosis might need to induce functional immune responses against OSP and Ipa, and that in endemic zones immune responses targeting OSP and active at mucosal surfaces may be paramount.

## Methods

### Luminex

*Shigella*-specific antibody subclass/isotype and Fcγ-receptor (FcγR) binding levels were assessed using a 384-well based customized multiplexed Luminex assay, as previously described(*39*). IpaB (WRAIR), IpaC (WRAIR), IpaD (WRAIR), *S. flexneri* 2a 2457T LPS (WRAIR), *S. flexneri* 3a strain J17B LPS (WRAIR), *S. flexneri* 6 LPS (icddrb), *S. sonnei* Moseley LPS (WRAIR), *S. flexneri* 2a OSP, *S. flexneri* 3a OSP, and *S. sonnei* OSP were used to profile - specific humoral immune response. OSP was purified from *S. flexneri* 2a Sf2a260214_1 LPS (icddrb), *S. flexneri* 3a Sf3a050214_5 LPS (icddrb), and *S. sonnei* Moseley LPS by acid hydrolysis and size exclusion chromatography and conjugated to BSA (Kelly M et al., in process, Vaccine). Tetanus toxin and Ebola (CEFTA, Mabtech Inc) were used as a control. Protein antigens were coupled to magnetic Luminex beads (Luminex Corp) by carbodiimide-NHS ester-coupling (Thermo Fisher). OSP and LPS antigens were modified by 4-(4,6-dimethoxy[1,3,5]triazin-2-yl)-4-methyl-morpholinium and conjugated to Luminex Magplex carboxylated beads. Antigen-coupled microspheres were washed and incubated with plasma samples at an appropriate sample dilution (1:50 for Isotypes and 1:100 for all Fc-receptors) for 2 hours at 37°C in 384-well plates (Greiner Bio-One). Unbound antibodies were washed away, and antigen-bound antibodies were detected by using a PE-coupled detection antibody for each subclass and isotype (IgG1, IgG2, IgG3, IgA1, IgA2 and IgM; Southern Biotech), and Fcγ-receptors were fluorescently labeled with PE before addition to immune complexes (FcγR2A, FcγR2B, FcγR3A, FcγR3B, FcαR; Duke Protein Production facility). After one hour incubation, plates were washed, and flow cytometry was performed with an IQue (Intellicyt), and analysis was performed on IntelliCyt ForeCyt (v8.1). PE median fluorescent intensity (MFI) was reported as a readout for antigen--specific antibody titers.

### Data Pre-processing

The raw MFI was scaled by the log10 function and was then subtracted by the corresponding PBS values. The normalized MFI values were assigned to zero if they were negative.

### Statistics

Microsoft Excel was used to compile and annotate experimental data. Violin plots, bar graphs, and x-y plots were generated in Graph Pad Prism V.8. Statistical differences between two groups were calculated using a two-sided Mann–Whitney test or Wilcoxon test for paired comparisons. To compare multiple groups, a Kruskal–Wallis test was used followed by the Dunn’s method correcting for multiple comparisons in Graph Pad Prism V.8 (significance levels: *p□<□0.05, **p□<□0.01, ***p□<□0.001, ****p□≤□0.0001). Heat maps and correlation matrix were created using Morpheus (https://software.broadinstitute.org/morpheus) or R version 1.4.1106.

### Antibody-dependent monocyte and neutrophil phagocytosis (ADMP and ADNP)

ADMP and ADNP were conducted as previously described(*40*). IpaB was biotinylated using Sulfo-NHS-LC-LC biotin (Thermo Fisher) and coupled to fluorescent Neutravidin-conjugated beads (Thermo Fisher). *S. flexneri* 2a OSP was modified with DMTMM and coupled to carboxylated fluorescent beads (Thermo Fisher). To form immune complexes, a mix of both, or *S. flexneri* 2a OSP only antigen-coupled beads was incubated for 2 hours at 37°C with diluted samples (1:200) and then washed to remove unbound immunoglobulins. For ADMP, the immune complexes were incubated for 4 hours with fresh blood monocytes isolated with commercially available kit (StemCell) (1.25×10^5^ monocytes/mL) and for ADNP for 1 hour with fresh blood neutrophils isolated from healthy donors with commercially available kit (StemCell). Following the incubation, cells were fixed with 4% PFA. For ADNP, neutrophils were washed, stained for CD66b (Biolegend), and then fixed in 4% PFA. For ADMP, monocytes were washed, stained for CD14 (Biolegend), and then fixed in 4% PFA. Flow cytometry was performed to identify the percentage of cells that had phagocytosed beads as well as the number of beads that had been phagocytosed (phagocytosis score = % positive cells × Median Fluorescent Intensity of positive cells/10000). Flow cytometry was performed with an IQue (Intellicyt), and analysis was performed on IntelliCyt ForeCyt (v8.1) or using FlowJo V10.7.1.

### Antibody dependent Complement deposition (ADCD)

ADCD was conducted using a 384-well based customized multiplexed assay. Protein antigens were coupled to magnetic Luminex beads (Luminex Corp) by carbodiimide-NHS ester-coupling

(Thermo Fisher). OSP and LPS antigens were modified by 4-(4,6-dimethoxy[1,3,5]triazin-2-yl)-4-methyl-morpholinium and conjugated to Luminex Magplex carboxylated beads(*41*). To form immune complexes, a mix of four antigen-coupled beads was incubated for 2 hours at 37°C with diluted samples (1:10) and then washed to remove unbound immunoglobulins. Lyophilized guinea pig complement (Cedarlane) was resuspended according to manufacturer’s instructions and diluted in gelatin veronal buffer with calcium and magnesium (Boston BioProducts). Resuspend guinea pig complement was added to immune complexes and incubated for 20 minutes at 37°C. Post incubation, C3 was detected with Fluorescein-Conjugated Goat IgG Fraction to Guinea Pig Complement C3 (Mpbio).

### IgA depletion

IgA was depleted from human plasma samples using CaptureSelect™ IgA Affinity Matrix (Thermo Fisher). The capture matrix was washed three times with PBS and incubated over night with 1:5 diluted plasma samples in a low protein binding MultiScreen® filter plate (Millipore). Depleted plasma was recovered by centrifugation of the filter plate. For each sample non-depleted plasma was treated similarly but without affinity matrix.

### Secondary neutrophil assays

Neutrophils of healthy blood donors were isolated using EasySep™ Direct Human Neutrophil Isolation Kit (Stemcell Technologies). Neutrophils were stimulated with immune-complexes as described above and supernatants collected after 4 h. MPO, lactoferrin and MMP-9 were detected in undiluted supernatant using ELISA kits (abcam and R&D systems).

### Computational analysis

A supervised multivariate analysis method of Least Absolute Shrinkage and Selection Operator (LASSO) followed by Partial Least Squares Regression (PLSR) was used to identify key antibody features that contribute to variation in the disease severity. Prior to building the LASSO-PLSR model, all titer, FcR and ADCD measurements were log transformed, and all measurements were then z-scored. LASSO identified a minimal set of features that drives separation in samples of varying disease severity. LASSO selected features were used to build the PLSR model regressing against the disease severity score. The performance of the algorithm was evaluated with R^2^ and Q^2^ metrics. Features were ranked based on their Variable of Importance (VIP) score and the loadings of the latent variable 1 (LV1) was visualized in a bar graph, which captures the contribution of each feature to the variation in disease severity. These analyses were carried out using R package *glmnet* (v4.0.2)(*42*) and *ropls* (v1.20.0) (*43*). Co-correlate networks were constructed based on the pairwise correlation between the top predictive features selected and all measured biophysical and functional features. Only correlations with an absolute Spearman correlation coefficient greater than 0.7 and *p*-value lower than 0.01 after correction for multiple comparisons by Benjamini-Hochberg (BH) were shown. Networks were generated using R package *network* (v1.16.0) [Butts C (2015). network: Classes for Relational Data.].

## Ethics statement

Sera were collected in fully IRB-approved challenge and field studies and secondary use for our current analysis was approved by the Institutional Review Board of the Massachusetts General Hospital.

## Financial Support

This research was supported through programs funded by the National Institutes of Health, including the National Institute of Allergy and Infectious Diseases (AI155414 [ETR, TRB, FQ], the Fogarty International Center, Training Grant in Vaccine Development and Public Health (TW005572 [MK]), and Emerging Global Fellowship Award TW010362 [TRB], and the Intramural Research Program of the NIH and NIDDK (PX and PK). The funders had no role in study design, data collection and analysis, decision to publish, or preparation of the manuscript.

The authors would like to thank the technical staff at the Naval Medical Research Unit Six (NAMRU-6) in Lima, Peru, Dr. Ryan Maves and Dr. Franca Jones for support of sample collection and access to bacterial isolate information and K. Ross Turbyfill for antigens (LPS and Ipa proteins) from Walter Reed Army Institute of Research.

## Bibliography

1. K. L. Kotloff, J. P. Nataro, W. C. Blackwelder, D. Nasrin, T. H. Farag, S. Panchalingam, Y. Wu, S. O. Sow, D. Sur, R. F. Breiman, A. S. G. Faruque, A. K. M. Zaidi, D. Saha, P. L. Alonso, B. Tamboura, D. Sanogo, U. Onwuchekwa, B. Manna, T. Ramamurthy, S. Kanungo, J. B. Ochieng, R. Omore, J. O. Oundo, A. Hossain, S. K. Das, S. Ahmed, S. Qureshi, F. Quadri, R. A. Adegbola, M. Antonio, M. J. Hossain, A. Akinsola, I. Mandomando, T. Nhampossa, S. Acácio, K. Biswas, C. E. O’Reilly, E. D. Mintz, L. Y. Berkeley, K. Muhsen, H. Sommerfelt, R. M. Robins-Browne, M. M. Levine, Burden and aetiology of diarrhoeal disease in infants and young children in developing countries (the Global Enteric Multicenter Study, GEMS): a prospective, case-control study. Lancet. 382, 209–222 (2013).

2. K. L. Kotloff, J. A. Platts-Mills, D. Nasrin, A. Roose, W. C. Blackwelder, M. M. Levine, Global burden of diarrheal diseases among children in developing countries: Incidence, etiology, and insights from new molecular diagnostic techniques. Vaccine. 35, 6783–6789 (2017).

3. S. Babji, G. Kang, Rotavirus vaccination in developing countries. Curr. Opin. Virol. 2, 443–448 (2012).

4. K. M. Rahman, S. El Arifeen, K. Zaman, M. Rahman, R. Raqib, M. Yunus, N. Begum, M. S. Islam, B. M. Sohel, M. Rahman, M. Venkatesan, T. L. Hale, D. W. Isenbarger, P. J. Sansonetti, R. E. Black, A. H. Baqui, Safety, dose, immunogenicity, and transmissibility of an oral live attenuated Shigella flexneri 2a vaccine candidate (SC602) among healthy adults and school children in Matlab, Bangladesh. Vaccine. 29, 1347–1354 (2011).

5. E. Richie, N. H. Punjabi, Y. Sidharta, K. Peetosutan, M. Sukandar, S. S. Wasserman, M. Lesmana, F. Wangsasaputra, S. Pandam, M. M. Levine, P. O’Hanley, S. J. Cryz, C. H. Simanjuntak, Efficacy trial of single-dose live oral cholera vaccine CVD 103-HgR in North Jakarta, Indonesia, a cholera-endemic area. Vaccine. 18, 2399–2410 (2000).

6. K. L. Kotloff, The Burden and Etiology of Diarrheal Illness in Developing Countries. Pediatr. Clin. North Am. 64, 799–814 (2017).

7. C. D, G. MS, B. C, R. T, O. I, Serum antibodies to lipopolysaccharide and natural immunity to shigellosis in an Israeli military population. J. Infect. Dis. 157, 1068–1071 (1988).

8. D. Cohen, M. S. Green, C. Block, R. Slepon, I. Ofek, Prospective study of the association between serum antibodies to lipopolysaccharide O antigen and the attack rate of shigellosis. J. Clin. Microbiol. 29, 386 (1991).

9. C. D, M.-S. S, B. A, A. V, G. S, A.-C. O, R. A, H. A, A. S, Serum IgG antibodies to Shigella lipopolysaccharide antigens - a correlate of protection against shigellosis. Hum. Vaccin. Immunother. 15, 1401–1408 (2019).

10. V. Rasolofo-Razanamparany, A. M. Cassel-Beraud, J. Roux, P. J. Sansonetti, A. Phalipon, Predominance of serotype-specific mucosal antibody response in Shigella flexneri-infected humans living in an area of endemicity. Infect. Immun. 69, 5230–5234 (2001).

11. M. M. Levine, K. L. Kotloff, E. M. Barry, M. F. Pasetti, M. B. Sztein, Clinical trials of Shigella vaccines: two steps forward and one step back on a long, hard road. Nat. Rev. Microbiol. 2007 57. 5, 540–553 (2007).

12. B. Bernshtein, E. Ndungo, D. Cizmeci, P. Xu, P. Kováč, M. Kelly, D. Islam, E. T. Ryan, K. L. Kotloff, M. F. Pasetti, G. Alter, Systems approach to define humoral correlates of immunity to Shigella. Cell Rep. 40, 111216 (2022).

13. K. A. Clarkson, K. R. Talaat, C. Alaimo, P. Martin, A. L. Bourgeois, A. Dreyer, C. K. Porter, S. Chakraborty, J. Brubaker, D. Elwood, R. Frölich, B. DeNearing, H. P. Weerts, B. Feijoo, J. Halpern, D. Sack, M. S. Riddle, V. G. Fonck, R. W. Kaminski, Immune response characterization in a human challenge study with a Shigella flexneri 2a bioconjugate vaccine. EBioMedicine. 66 (2021), doi:10.1016/J.EBIOM.2021.103308.

14. R. A. Oberhelman, D. J. Kopecko, E. Salazar-Lindo, E. Gotuzzo, J. M. Buysse, M. M. Venkatesan, A. Yi, C. Fernandez-Prada, M. Guzman, R. Leon-Barua, R. B. Sack, Prospective study of systemic and mucosal immune responses in dysenteric patients to specific Shigella invasion plasmid antigens and lipopolysaccharides. Infect. Immun. 59, 2341–2350 (1991).

15. F. R. Jones, J. L. Sanchez, R. Meza, T. M. Batsel, R. Burga, E. Canal, K. Block, J. Perez, C. T. Bautista, J. Escobedo, S. E. Walz, Short report: High incidence of Shigellosis among Peruvian soldiers deployed in the Amazon river basin. Am. J. Trop. Med. Hyg. 70, 663–665 (2004).

16. R. W. Frenck, M. Dickey, A. E. Suvarnapunya, L. Chandrasekaran, R. W. Kaminski, K. A. Clarkson, M. McNeal, A. Lynen, S. Parker, A. Hoeper, S. Mani, A. Fix, N. Maier, M. M. Venkatesan, C. K. Porter, Establishment of a Controlled Human Infection Model with a Lyophilized Strain of Shigella sonnei 53G. mSphere. 5 (2020), doi:10.1128/MSPHERE.00416-20.

17. G. Robin, D. Cohen, N. Orr, I. Markus, R. Slepon, S. Ashkenazi, Y. Keisari, Characterization and quantitative analysis of serum IgG class and subclass response to Shigella sonnei and Shigella flexneri 2a lipopolysaccharide following natural Shigella infection. J. Infect. Dis. 175, 1128–1133 (1997).

18. M. H. Nahm, J. Yu, H. P. Weerts, H. Wenzel, C. S. Tamilselvi, L. Chandrasekaran, M. F. Pasetti, S. Mani, R. W. Kaminski, Development, Interlaboratory Evaluations, and Application of a Simple, High-Throughput Shigella Serum Bactericidal Assay. mSphere. 3 (2018), doi:10.1128/msphere.00146-18.

19. F. Nimmerjahn, J. V. Ravetch, Fc-Receptors as Regulators of Immunity. Adv. Immunol. 96, 179–204 (2007).

20. M. Van Egmond, C. A. Damen, A. B. Van Spriel, G. Vidarsson, E. Van Garderen, J. G. J. Van De Winkel, IgA and the IgA Fc receptor. Trends Immunol. 22, 205–211 (2001).

21. D. Cohen, C. Block, M. S. Green, G. Lowell, I. Ofek, Immunoglobulin M, A, and G antibody response to lipopolysaccharide O antigen in symptomatic and asymptomatic Shigella infections. J. Clin. Microbiol. 27, 162–167 (1989).

22. C. D, G. MS, B. C, S. R, O. I, Prospective study of the association between serum antibodies to lipopolysaccharide O antigen and the attack rate of shigellosis. J. Clin. Microbiol. 29, 386–389 (1991).

23. K. Nothelfer, E. T. Arena, L. Pinaud, M. Neunlist, B. Mozeleski, I. Belotserkovsky, C. Parsot, P. Dinadayala, A. Burger-Kentischer, R. Raqib, P. J. Sansonetti, A. Phalipon, B lymphocytes undergo TLR2-dependent apoptosis upon Shigella infection. J. Exp. Med. 211, 1215–1229 (2014).

24. G. Sellge, J. G. Magalhaes, C. Konradt, J. H. Fritz, W. Salgado-Pabon, G. Eberl, A. Bandeira, J. P. Di Santo, P. J. Sansonetti, A. Phalipon, Th17 cells are the dominant T cell subtype primed by Shigella flexneri mediating protective immunity. J. Immunol. 184, 2076–2085 (2010).

25. H. Ashida, M. Ogawa, M. Kim, S. Suzuki, T. Sanada, C. Punginelli, H. Mimuro, C. Sasakawa, Shigella deploy multiple countermeasures against host innate immune responses. Curr. Opin. Microbiol. 14, 16–23 (2011).

26. D. Islam, B. Christensson, Disease-dependent changes in T-cell populations in patients with shigellosis. APMIS. 108, 251–260 (2000).

27. R. Hamre, I. N. Farstad, P. Brandtzaeg, H. C. Morton, Expression and Modulation of the Human Immunoglobulin A Fc Receptor (CD89) and the FcR γ Chain on Myeloid Cells in Blood and Tissue. Scand. J. Immunol. 57, 506–516 (2003).

28. F. Geissmann, P. Launay, B. Pasquier, Y. Lepelletier, M. Leborgne, A. Lehuen, N. Brousse, R. C. Monteiro, A subset of human dendritic cells expresses IgA Fc receptor (CD89), which mediates internalization and activation upon cross-linking by IgA complexes. J. Immunol. 166, 346–352 (2001).

29. A. Gorter, P. S. Hiemstra, P. C. Leijh, M. E. van der Sluys, M. T. van den Barselaar, L. A. van Es, M. R. Daha, IgA- and secretory IgA-opsonized S. aureus induce a respiratory burst and phagocytosis by polymorphonuclear leucocytes. Immunology. 61, 303 (1987).

30. E. Aleyd, M. W. M. van Hout, S. H. Ganzevles, K. A. Hoeben, V. Everts, J. E. Bakema, M. van Egmond, IgA enhances NETosis and release of neutrophil extracellular traps by polymorphonuclear cells via Fcα receptor I. J. Immunol. 192, 2374–2383 (2014).

31. I. S. Hansen, L. Krabbendam, J. H. Bernink, F. Loayza-Puch, W. Hoepel, J. A. Van Burgsteden, E. C. Kuijper, C. J. Buskens, W. A. Bemelman, S. A. J. Zaat, R. Agami, G. Vidarsson, G. R. Van Den Brink, E. C. De Jong, M. E. Wildenberg, D. L. P. Baeten, B. Everts, J. Den Dunnen, FcαRI co-stimulation converts human intestinal CD103+ dendritic cells into pro-inflammatory cells through glycolytic reprogramming. Nat. Commun. 9 (2018), doi:10.1038/S41467-018-03318-5.

32. R. Raqib, A. A. Lindberg, B. Wretlind, P. K. Bardhan, U. Andersson, J. Andersson, Persistence of local cytokine production in shigellosis in acute and convalescent stages. Infect. Immun. 63, 289–296 (1995).

33. R. Raqib, Å. Ljungdahl, A. A. Lindberg, U. Andersson, J. Andersson, Local entrapment of interferon gamma in the recovery from Shigella dysenteriae type 1 infection. Gut. 38, 328–336 (1996).

34. M. M. Gomes, S. B. Wall, K. Takahashi, J. Novak, M. B. Renfrow, A. B. Herr, Analysis of IgA1 N-glycosylation and its contribution to FcαRI binding. Biochemistry. 47, 11285– 11299 (2008).

35. T. S. Mattu, R. J. Pleass, A. C. Willis, M. Kilian, M. R. Wormald, A. C. Lellouch, P. M. Rudd, J. M. Woof, R. A. Dwek, The glycosylation and structure of human serum IgA1, Fab, and Fc regions and the role of N-glycosylation on Fcα receptor interactions. J. Biol. Chem. 273, 2260–2272 (1998).

36. U. Steffen, C. A. Koeleman, M. V. Sokolova, H. Bang, A. Kleyer, J. Rech, H. Unterweger, M. Schicht, F. Garreis, J. Hahn, F. T. Andes, F. Hartmann, M. Hahn, A. Mahajan, F. Paulsen, M. Hoffmann, G. Lochnit, L. E. Muñoz, M. Wuhrer, D. Falck, M. Herrmann, G. Schett, IgA subclasses have different effector functions associated with distinct glycosylation profiles. Nat. Commun. 2020 111. 11, 1–12 (2020).

37. E. Ndungo, L. R. Andronescu, A. G. Buchwald, P. Mawindo, K. Laufer, M. F. Pasetti, Repertoire of naturally acquired maternal antibodies transferred to infants for 1 protection against shigellosis 2 3, doi:10.1101/2021.05.21.445178.

38. E. Ndungo, J. B. Holm, S. Gama, A. G. Buchwald, S. M. Tennant, M. K. Laufer, M. F. Pasetti, D. A. Rasko, Dynamics of the Gut Microbiome in Shigella-Infected Children during the First Two Years of Life. mSystems. 7 (2022), doi:10.1128/MSYSTEMS.00442-22/SUPPL_FILE/MSYSTEMS.00442-22-S0005.XLSX.

39. B. EP, W. JA, L. S, N. H, N. E, B. DH, A. G, S.-K. M, A. ME, Optimization and qualification of an Fc Array assay for assessments of antibodies against HIV-1/SIV. J. Immunol. Methods. 455, 24–33 (2018).

40. A. L. Butler, J. K. Fallon, G. Alter, A sample-sparing multiplexed ADCP assay. Front. Immunol. 10, 1851 (2019).

41. S. A. Schlottmann, N. Jain, N. Chirmule, M. T. Esser, A novel chemistry for conjugating pneumococcal polysaccharides to Luminex microspheres. J. Immunol. Methods. 309, 75– 85 (2006).

42. J. Friedman, T. Hastie, R. Tibshirani, Regularization Paths for Generalized Linear Models via Coordinate Descent. J. Stat. Softw. 33, 1 (2010).

43. E. A. Thévenot, A. Roux, Y. Xu, E. Ezan, C. Junot, Analysis of the Human Adult Urinary Metabolome Variations with Age, Body Mass Index, and Gender by Implementing a Comprehensive Workflow for Univariate and OPLS Statistical Analyses. J. Proteome Res. 14, 3322–3335 (2015).

